# From intent to implementation: Factors affecting public involvement in life science research

**DOI:** 10.1101/748889

**Authors:** John. A. Burns, Kora Korzec, Emma R. Dorris

## Abstract

Public involvement is key to closing the gap between research production and research use, and the only way to achieving ultimate transparency in science. The majority of life science research is not public-facing, but is funded by the public and impacts the community. We undertook a survey of researchers within the life sciences to better understand their views and perceived challenges to involving the public in their research. We had a valid response cohort of n=110 researchers, of whom 90% were primarily laboratory based. Using a mixed methods approach, we demonstrate that a top-down approach is key to motivate progression of life scientists from feeling positive towards public involvement to actually engaging in it. Researchers who viewed public involvement as beneficial to their research were more likely to have direct experience of doing it. We demonstrate that the systemic flaws in the way life sciences research enterprise is organised, including the promotion system, hypercompetition, and time pressures are major barriers to involving the public in the scientific process. Scientists are also apprehensive of being involuntarily involved in the current politicized climate, misinformation and publicity hype surrounding science nowadays makes them hesitant to share their early and in-progress research. The time required to deliberate study design and relevance, plan and build relationships for sustained involvement, provide and undertake training, and improve communication in the current research environment is often considered nonpragmatic, particularly for early career researchers. In conclusion, a top-down approach involving institutional incentives and infrastructure appears most effective at transitioning researchers from feeling positive towards public involvement to actually implementing it.

## Introduction

Scientific knowledge is a public good[1,2], not solely due to its intrinsic properties but also as a source of diversity, flexibility and innovation[1,3]. In line with the increase in citizen science and patient and public involvement (PPI) initiatives, there is increasing focus on the translation and implementation of new scientific evidence in real-world settings[4]. Across the life sciences, narrowing the gap between research production and research use is a key challenge. There is increasing evidence that public involvement and stakeholder engagement is key to achieving impact and true provision and use of scientific knowledge for the benefit of society[5–11].

Scientific questions range in scale from the subatomic through questions encompassing the entire universe [12,13]. Data collection and interpretation from projects that span large geographic areas or require human curation of extremely large datasets can fall beyond the scope of an individual research lab. When this is the case, scientists often recruit and train non-specialists to participate in aspects of these grand projects. Projects in this realm include an ongoing census of all birds[14] or butterflies[15] across a continent, monitoring coral reefs for bleaching[16], and monitoring sea turtle populations[17] among many others. Research projects that include members of the public as active participants are called “citizen science” efforts.

Citizen science efforts where members of the public work with professional scientists can be classified into three major categories: contributory, collaborative, and co-created projects[18]. The categories reflect the level of input and participation from members of the public. Projects like Galaxy Zoo[13], Cornell Ornithology Lab[19], and Foldit[20] are contributory projects where members of the public are gathering data for a project designed, managed, and interpreted by research scientists. Collaborative projects include community monitoring efforts, such as water quality monitoring[21] and local conservation efforts[22]. Collaborative projects are often designed by scientists but receive input and feedback from participants and legislators to achieve a conservation goal, such as maintaining water quality or protecting habitat for a threatened species[21,22]. Co-created projects occur when members of the public have a say in study design, data collection, data interpretation, and dissemination of results[18]. Co-created projects can help frame concerns of community members as scientific questions and can levy specialized local knowledge and engagement from community members. One such co-created project, “Gardenroots” involved measuring arsenic exposure routes in a community based near a superfund site in Arizona[23]. Involving members of the community in all aspects of the study aided the overall goal of risk communication to the community since members of the community were involved in the actual risk analysis [23].

Apart from expanding the reach of small research groups to larger geographic areas and datasets, citizen science efforts often have goals of outreach and education. Engaging the public in scientific pursuits so they are more informed about how science works, what a research project entails, and what are realistic expectations and limitations of typical research projects[24,25]. A key aspect of citizen science efforts is training volunteers, who are typically not experts in the scientific field of study, for accurate and reproducible data collection[26]. One way to oversee volunteer data collection is to validate a subset of the observations by coincident or re-analysis by an expert and to develop and implement tools for data sharing and evaluation[26,27]. Citizen science programs, unlike a typical laboratory experiment, are often survey efforts and are not designed to test a specific hypothesis[25]. Building a project around a specific question can help focus efforts and guide outcomes. Successful completion of projects that include a citizen science component is aided by careful and deliberate study design including project goals, scientific questions, plans for sustained participation, training, communication, and desired project outcomes[28].

Patient and public involvement (PPI) in health research is akin to citizen science applied specifically to the health and social care field. The most commonly used PPI definition is “research carried out ‘with’ or ‘by’ members of the public rather than ‘to’, ‘about’ or ‘for’ them”[29]. Both citizen science and PPI are most commonly practiced by public-facing disciplines. Expanding public involvement to laboratory-based and non-public facing disciplines is more challenging for all stakeholders [30,31]. In this study we use an international survey of researchers within the life sciences as a method of understanding the attitudes and challenges of public involvement and to identify the factors that promote public involvement in life science research.

#### Box 1. Definitions used during this study

**Participation:** public take part in an experiment, trial, or study as a subject. The person who participates is the target of observation by researchers. They are data providers. Examples: participant in a clinical trial.

**Engagement:** a one-way dialogue from researcher to public where the researcher explains/educates/informs the public about their research. Examples: public lectures, lab tours, research demonstrations

**Consultation:** a one-way dialogue from public to researcher where the public's input on matters affecting them is sought. The public are usually not decision makers. Examples: Surveys or focus groups to inform policy or governance

**Involvement:** two-way dialogue between public and researchers. The active involvement between people who use services, the general public and researchers. Public are active collaborators, akin to researchers, clinicians/professionals or managers who are asked for their views and experiences when contributing to research. The public usually have some degree of decision making. Examples: Research advisory groups members, citizen science, public educators or mentors.

## Materials and Methods

This international, voluntary, quantitative study used convenience sampling to collect data from an anonymous survey administered to self-reported life scientists.

### Participants

Surveys were distributed electronically to members of the international working group of life scientists, the eLife Ambassadors for Good Practice in science (n=206) via a private online cloud-based team collaboration hub and via email, with a request for them to also share it within their scientific communities, societies within the life sciences, and through internal institutional communications. We received 122 responses, of which 117 individuals self-reported as life scientists. A further five respondents did not consent to the use of their data, and two were under the age of 18, resulting in a valid response cohort of n=110.

### Public Involvement in Life Sciences Survey

The survey consisted of five sections: 1. Consent; 2. Research Details; 3. Researcher Demographics; 4. Public Involvement Awareness and Experience; and 5. Views on Public Involvement. Sections 1-4 contained nominal questions whereas section five contained both nominal and unrestricted textbox questions.

### Analysis

Data was analysed in IBM SPSS v24. Factor analysis with correlation matrix was used to identify co-linearity and redundancy within the variables. Nominal variables were analysed using Pearson’s Chi-squared test or Fisher’s exact test, using an alpha level of 0.05. Significance testing related to public involvement experience used 1-tailed tests. Free text answers underwent qualitative textual analysis. Data were analysed using the pragmatist approach of inductive thematic analysis [32] combined with frequency analysis of themes, and managed using excel [33]. Two researchers (KK; ED) analysed data independently, with a third independent researcher nominated to mediate potential discrepancies.

## Results

### Research Demographics

In the total valid response cohort of 110 researchers, 23 different areas of the life sciences were identified as primary research areas (figure 1A); with a median of 5 respondents per research area (range 1-20). Respondents were overwhelmingly laboratory-based at a university or research institute (n=99 (90.8%); Fig 1B).

**Figure 1:**
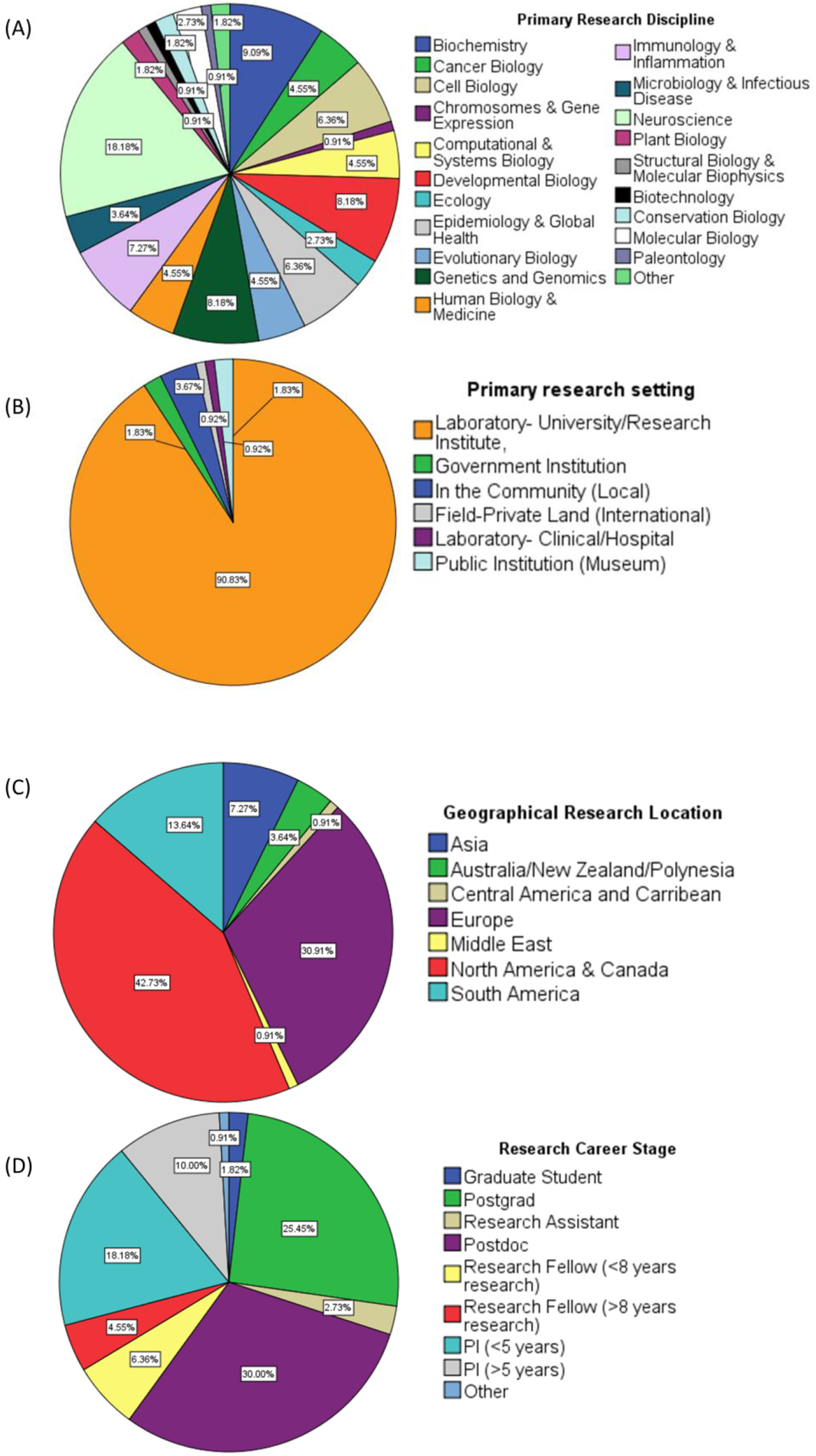
Researcher Demographics. (A) Primary research discipline of responders (B) primary research setting (C) Geographical location (D) Career Stage.

Respondents were based in research locations across five continents (Fig 1C) with no respondents from research institutes in Africa. The number of respondents from Asian research institutes was relatively low considering the high density of researchers in Asia (Asia and the Middle East based responders n=9 (8.2% of all responses))[34]. However, it should be noted that the survey was only available in the English language. North American and European-based research (n=47 (42.8% of all responses) and n=34 (30.9% of all responses) respectively) followed by Central and South American-based researchers (n=16 (14.4%). Oceania -based researchers accounted for 3.6% of respondents (n=4). No association was found between geographical research location and experience of public involvement (Fisher’s exact Chi-square 8.803, p=0.132).

The research experience/career level of respondents ranged from graduate student (1.8%) to principal investigator (28.18%; Fig 1D). 30% of respondents were postdocs, and 10.9% Research Fellows. Experience of public involvement did not vary by career stage (Pearson’s Chi-squared 1.624, p=0.654).

### Researcher Demographics and Public Involvement

The likelihood of having involved the public in research did not associate with researcher’s sex (p=0.112). 42.7% (n=47) of respondents were female and 55.5% (n=61) were male; 27.65% of female and 32.79% of male researchers had involved the public in their research at least once. We tested six additional demographic variables for association with having involved the public in research (Table 1). Age positively associated with experience of public involvement (correlation coefficient 0.323; linear-by-linear association statistic 6.501, p=0.006 (Exact 1-sided), Fig. 2A). This is in agreement with the literature demonstrating that both scientific engagement and active dissemination increase with age and experience[35,36]

**Table 1:** Association of researcher demographics with public involvement experience.

| Question | Correlation<br>with PI | p-value<br>(1-tailed) |
| --- | --- | --- |
| What sex are you? | -0.015 | 0.442 |
| What age are you? | 0.323 | 0.001 |
| What nationality are you? | -0.026 | 0.400 |
| What is your Ethnic/Cultural Background? | 0.158 | 0.059 |
| Socio-economic background: At what age did your parent/guardian leave continuous full-time education? | 0.193 | 0.028 |
| Do you have a disability? | 0.066 | 0.258 |
| Have you applied for funding or ethics where there was a specific question on public involvement in research? | 0.437 | 0.000 |
| Have you involved the public in your research? | 1.000 |  |

**Fig. 2:**
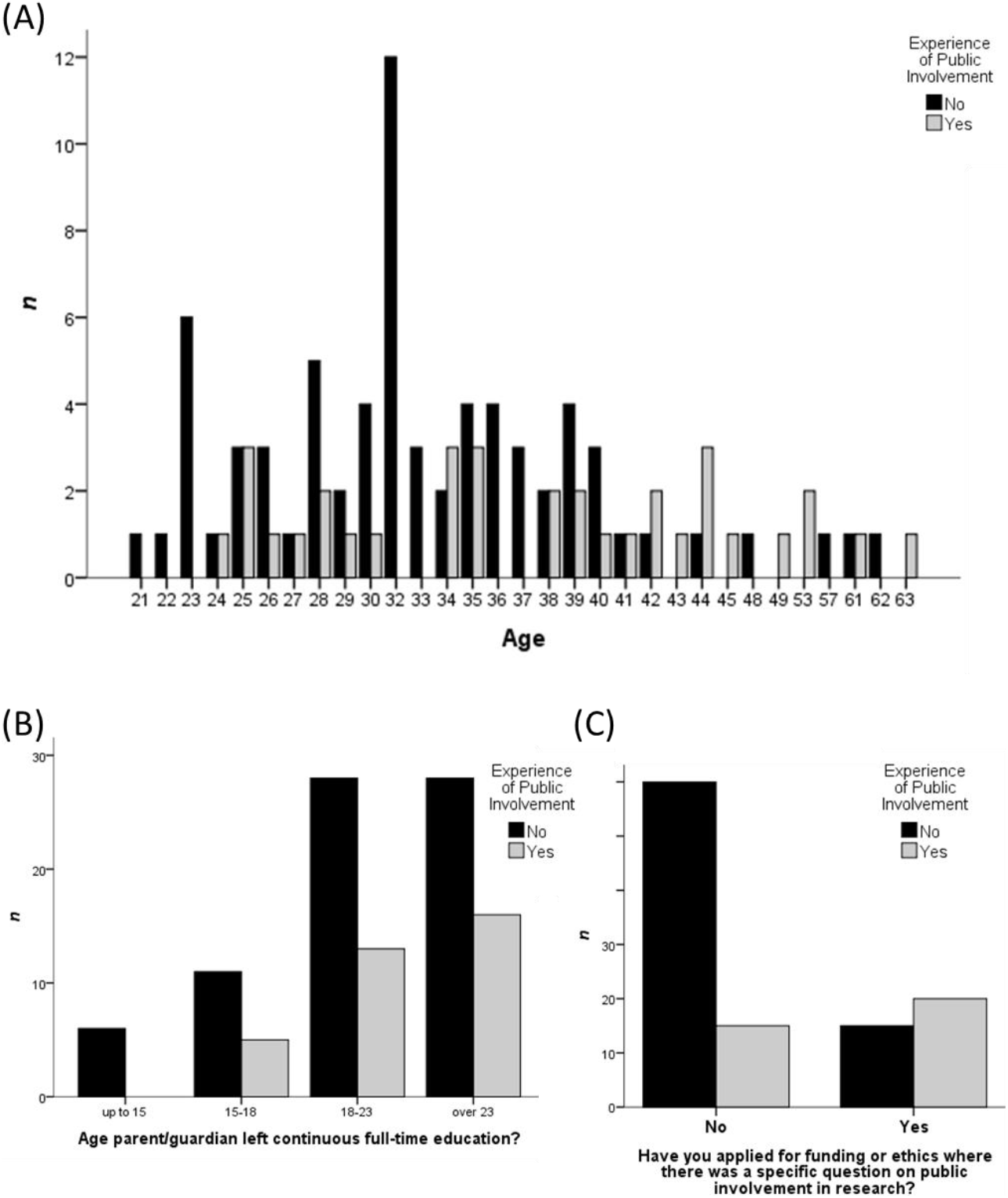
Factors correlating with public involvement. Age has a positive correlation with public involvement experience(A), as does the (B) age at which a researchers parent left full time education and (C) whether a researcher has been asked a specific question on public involvement in a funding or ethics application or not.

A single question on the age that parents finished education was used as a proxy for researcher socio-economic status (SES) during upbringing[37]. A positive correlation (correlation coefficient 0.193, p=0.028) was observed between the age at which a researcher’s parent/guardian left full time education and public involvement (Fig. 2B). However, Chi-square test for linear by linear association was not significant at the α0.05 level (Chi statistic 1.960, p=0.099 (Exact 1-sided)).

### Agency and institutional policies can promote public involvement

The largest fraction of variance in public involvement was explained by whether a researcher had applied for funding or had ethics submissions where there was a specific question on public involvement in research. This variable accounted for 22.9% of the variance in our data (principal component analysis) and correlated, with public involvement via factor loading (promax rotation with Kaiser normalisation). A positive correlation was observed between having been asked a specific question on public involvement in an ethics or funding application, and experience of public involvement (Table 2; correlation coefficient 0.437; p<0.000) attributed to a linear relationship between the variables (linear by linear association statistic 15.038; p=0.000).

**Table 2:** Awareness of public involvement and correlation with public involvement experience

| **Question** | **No** | **Maybe** | **Yes** | **Correlation**<br>**with PI** | **p-value**<br>**(1-**<br>**tailed)** |
| --- | --- | --- | --- | --- | --- |
| Would you like to involve the public in your research? | 5<br>(4.55%) | 47<br>(42.73%) | 58<br>(52.73%) | 0.149 | 0.060 |
| As far as you know, does your research institute provide resources (excluding funding) for the involvement of the public in research? | 39<br>(35.54%) | 16<br>(14.55%) | 55<br>(50%) | 0.007 | 0.470 |
| As far as you know, does your research institute encourage the involvement of the public in research? | 25<br>(22.73%) | n/a | 85<br>(77.27%) | -0.049 | 0.307 |
| Have you completed any training to facilitate the involvement of the public in your research? | 87<br>(79.09%) | n/a | 23<br>(20.91%) | 0.321 | 0.000 |
| Are you aware of online resources to help facilitate public involvement in research? | 75<br>(68.18%) | n/a | 35<br>(31.82%) | 0.204 | 0.016 |
| Have you used any online resources for the involvement of the public in research? | 89<br>(80.91%) | n/a | 21<br>(19.09%) | 0.463 | 0.000 |
| Have you been offered training to facilitate the involvement of public in your research? | 78<br>(70.91%) | n/a | 32<br>(29.09%) | 0.293 | 0.001 |
| Do you think public involvement in research is beneficial to the general public? | 1<br>(0.91%) | 12<br>(10.91%) | 97<br>(88.18%) | 0.079 | 0.206 |
| Do you think public involvement in research is beneficial to research? | 1<br>(0.91%) | 35<br>(31.82%) | 74<br>(51.82%) | 0.229 | 0.008 |
| Do you think public involvement in research is beneficial to you as a researcher? | 10<br>(9.09%) | 43<br>(39.09%) | 57<br>(51.82%) | 0.270 | 0.002 |
| Do you think you think some level of public involvement should be a requisite/part of the terms of funding for all research projects? | 44<br>(40%) | 9<br>(8.18%) | 16<br>(14.55%) | 0.125 | 0.096 |
|  |  | 24*<br>(21.82%) | 17*<br>(15.45%) |  |  |
| Have you involved the public in your research? | 75 | n/a | 35 | 1.000 |  |

\* Only for publicly funded research

Public involvement experience was correlated with training in public involvement (correlation coefficient 0.321, p=0.000); and both awareness of and use of online public involvement resources (0.204, p=0.016; 0.463, p=0.000, respectively).

### Researchers involve the public when they consider such involvement as beneficial to themselves or to their research

Respondents overwhelmingly agreed that public involvement in life science research was beneficial to the general public. This did not correlate with experience of public involvement; however only one respondent considering it not to be beneficial (0.91%); 10.91% (n= 12) maybe; and 88.18% (n=97) considered public involvement beneficial to the general public (Table 2). There was a positive correlation between researchers who considered public involvement beneficial to the researcher themselves (0.270, p=0.002) or to their research (0.229, p=0.008).

### The academic career track is a barrier to public involvement

As per Maccarthy et al (2019), challenges surrounding public involvement were asked in the context of institutional barriers, personal worries and research concerns[38]. There were 81 responses to the question on institutional barriers [In your view, what are the institutional barriers to including public involvement in your research?]. The well documented challenges of regulations, resources and funding were major themes emerging from the data [10,38–40]. The other major theme was that of career benefit.

> *“Not valued as academic currency for TT*/jobs.”*
>
> — - Postdoctoral researcher in cell biology, based in North America (*TT: Tenure Track)

Public involvement is demanding both in terms of time and resources. The academic career structure and the increasingly high bar to earn job stability or tenure as an academic researcher is a major barrier to public involvement[41,42]. This is reflected in our survey data, where we show merely feeling positively towards public involvement did not associate with implementing involvement in research; and the likelihood of having involved the public in research increased with age.

> *“The extremely competitive science system preclude me to really consider it”*
>
> — - Research Fellow in biochemistry, based in Europe.

Institutions need to consider not only the funding and resources available for public involvement, but the environment necessary to promote it. The continued focus on academic productivity, at the expense of academic activities that grow or improve research, will continue to restrict researchers in their innovative potential and growth as pragmatism and limited time and energy necessitates focus on tenure track and promotion variables. The attitude of “waiting for tenure to do what you actually love” has become pervasive in the life sciences. However, many excellent and innovative scientists are lost to science during the long road to tenure [43–45]. Engaging stakeholders is beneficial to science and to the translation and use of scientific knowledge[5,9,46] and institutions need to consider the life sciences system as a whole in their efforts to encourage stakeholder engagement and knowledge transfer[45].

### Fear of misrepresentation and public involvement

We had 79 responses to the question on personal barriers to involvement, of which n=9 indicated they did not have any personal barriers. Within the remaining responses to the question [In your view, what are the personal barriers (worries) that would prevent you including public involvement in your research?] two main, and highly related, themes emerged: fear of misrepresentation and personal communication skills.

Science communication is a critical skill for public involvement. Without some grasp of the relevant science, the public cannot be expected to make informed decisions about research[47]. In the absence of a pre-existing relationship, how can a researcher understand the public’s information needs, and make it accessible in a format useful for public involvement? Can we expect laboratory-based life scientists to also be skilled in bridging science and decision making? There are frameworks available to assist in science communication but the evidence-base for their effectiveness in the life sciences is understudied [48–50].

> *“Once misinformation has been disseminated, it seems very difficult to rectify in the current information sharing climate”*
>
> — - Postdoc in Neuroscience, based in Europe

Science has become increasingly politicised in the public sphere[51]. Political orientations and ideologies can shape public trust in science[51,52]. Fear of misrepresentation of their research, either intentional or otherwise, was a major theme identified in our data.

> *“There is always a fear that your words might be taken out of context to push an agenda that you don’t agree with. Also, putting yourself out there may make you vulnerable to verbal or even physical attacks on you”*
>
> — - Principal Investigator in Biochemistry, based in Europe

Complicating the issue is the reach of misinformation. Once misinformation has entered the public sphere, rebuttal with corrections to the misinformation tend to be muted compared with the propagation of the misinformation, leading to false and misrepresented information remaining within the pool of common knowledge [53]. Our data shows this has made researchers wary of opening their in-progress research to the public, even when they recognise there are potential benefits to their research of doing so.

Politicized environments often induce suspicions about science communicators’ true motives or expertise and questions may arise as to whether a scientist can really be trusted[54]. There is a fear among scientists about the potential personal backlash stemming from public involvement. This links to the theme of career benefit.

> *“Fear of being misrepresented or being “less serious””*
>
> — - Postgraduate conservation biologist, South America

Involving the public leaves you and your research vulnerable to misrepresentation and being negatively viewed by your peers and colleagues. Public involvement may also raise a scientists’ public profile, leading to backlash. Known as “The Sagan Effect”[55], this is the concept that public engagement can be a detriment to an academic’s career and stems from a common, but erroneous perception in the sciences that scientists who perform outreach activities are less successful academically[35,36,56]. This causes dissonance between the perceived reduced credibility of those who engage the public and the underpinning belief that public involvement is beneficial and increasingly necessary to meet their duties of knowledge transfer as well as knowledge generation[57].

### Is public involvement an efficient use of time?

When asked expressly about the concerns of public involvement to their research [In your view, what are the research concerns that would prevent you including public involvement in your research?]; of the 70 respondents, 15 said they had no concerns about its impact on their research. Of the remaining respondents: misrepresentation, workload, effect on research focus and relevance of public involvement for their research emerged as concerns with approximately equal frequency.

> *“The main barrier is time - being involved in public outreach takes precious time away from research activities, getting grants and publications.”*
>
> — - Principal Investigator, cancer and cell biology, Europe.

Time is a precious resource in the life sciences and burnout is common [58–61]. Even when researchers acknowledged the potential benefit of public involvement to their research, there was concern that there were not sufficient resources in place such that the benefit to their research would not outweigh the cost in terms of time.

## Discussion

Science is a public good and a social enterprise. In addition to inordinate technological development, the rapid growth in life sciences research and knowledge has occurred due to a long-standing public investment in life science research [45,62–64]. The fact that much research is not public-facing does not mean that it is not publicly relevant. In order to continue sustained investment into life science research, we must build more comprehensive, interactive and mutually beneficial opportunities for dialogue and exchange with our largest funders- the public[65]. Openness in science does not solely relate to article processing fees[66]. Responsibility and openness in science also relates to increasing the accessibility and use of scientific knowledge. To this end, responsible science should include appropriate dialogue with the community upon which it has an impact; or which it may be perceived to impact [67,68].

Here we demonstrate that a top-down approach, understood as funders and institutions providing the mandate and resources that support it, encourages public involvement in life sciences research. Simply feeling positively towards involving the public in research was not sufficient for researchers to implement it. Researchers who considered public involvement to be beneficial to the researcher themselves or to their research, rather than just to the general public, were more likely to engage in it. This suggests that provision of greater context or immersive experience of public involvement in the life sciences would allow researchers to understand and conceptualise the potential benefit of public involvement to their own research.

In line with previous findings from public engagement and public communications, the likelihood of having involved the pubic in research increased with age[35,36]. Our qualitative analysis identified that the career pathway, including funding pressures and the promotion system of research institutions has a key role to play in the pragmatic ability of early-career researchers to truly consider public involvement. This highlights that simply providing training and resources for public involvement, without implementing changes in the science career pathways, may have limited benefit at an early career stage[69].

Researchers who had involved the public in their research were more knowledgeable about the local and electronic resources available to them and more likely to have engaged in training for public involvement. These resources seem helpful downstream, once public involvement has been initiated.

Researcher vulnerability is an important issue often overlooked in public involvement. For institutions wishing to encourage increased public involvement they may need to consider what protections are in place for researchers exposing their early stage and in-development research to the public sphere. The time required to deliberate study design and relevance, plan and relationship build for sustained involvement, provide and undertake training, and improve communication in the current research environment is not considered feasible within their research and/or institutional environment for many researchers and is an area in need of further research into the best ways to overcome these challenges. Misrepresentation (intentional or otherwise) coupled with the ecosystem of rapid spread of false information on public platforms, and the fear thereof, is a complex issue[70]. It is made more complex in the context of non-public interfacing research, which does not have a high level of public understanding in the first place [71–73]. To truly integrate public involvement in a mutually beneficial way in non-public-facing disciplines requires deeper understanding and improved holistic support for this already over-burdened researcher demographic[45].

This study highlight the challenges faced in compelling researchers, particularly early-career laboratory science researchers, to engage directly with the public by involving them in their research projects. Despite the challenges highlighted here, citizen science efforts have been growing in terms of number of projects[74], outcomes in terms of publications[74], and the number or resources available to researchers and members of the public alike including open source, peer reviewed journals on citizen science and public and patient involvement [75–78]. Our results emphasize the need for integration and incentivization of the use of these resources by institutions and funding agencies to maximize researcher’s interests in public involvement[69]. Effective change towards a more open and inclusive model of life sciences will require more than just training and funding. Policy makers and institutions can greatly influence the implementation of public involvement in research through the working conditions and environment they promote.

## Limitations

The sample size of 110 researchers is small, limiting the capacity to conduct more in-depth statistical associations. The high rate of consensus on the beneficial nature of public involvement indicates selection bias in the respondents toward those who view public involvement positively. The survey was only available in English, limiting the potential respondents to those proficient in written English.

### Conclusions

Fundamentally, a researcher’s decision to involve the public will be decided by whether they believe it has sufficient potential value to both themselves and their research as to warrant the investment in terms of time, energy and potential or perceived career consequences. Policy makers and institutions can greatly influence this decision by creating an environment supportive of responsible research practices, including public involvement. Creating this environment in many cases will require a paradigm shift if public involvement is to become more than a marginal curiosity or tokenistic effort in laboratory-based research[79].

## Acknowledgements

This study was conducted as an initiative of the eLife Ambassadors for Good Practice in Science; we would like to acknowledge the help and support of the eLife Ambassadors and community members. In particular Dr. Erin Wissink, Dr. Hannah Wang and Dr. Vinodh Ilangovan.

